# Enabling Stable Cortical Reconstruction in the HCP Pipeline at Standard Resolution Using FastSurfer Integration with Hybrid T2w- and T1w-based Masking

**DOI:** 10.64898/2026.05.29.728714

**Authors:** Koji Hatano, Hirofumi Hirakawa, Hiroyuki Matsuda, Nobuhiko Hoaki, Takeshi Terao, Tsuyoshi Shimomura

## Abstract

This study aimed to enable stable cortical reconstruction within the Human Connectome Project (HCP) pipeline under standard-resolution (1 mm³) conditions by developing an integrated framework combining FastSurfer with a hybrid T2w- and T1w-based masking strategy. We implemented a FastSurfer-integrated HCP pipeline incorporating a hybrid masking approach (t2log-hybrid) to stabilize cortical reconstruction, where SynthStrip-derived masks were refined using log-transformed T2-weighted and squared T1-weighted images. An anterior commissure–anchored spatial switching mechanism selectively replaced artifact-prone T2w regions with T1w-derived information in orbitofrontal areas. Performance was evaluated against FreeSurfer v6 (FS6), v7 (FS7), and FastSurfer using geometric agreement, regional consistency, and vertex-wise thickness–myelin analyses. The proposed framework reduced total processing time by over 50% compared with FS6 and maintained high geometric agreement with modern pipelines (vertex-wise *r* = 0.970 vs FastSurfer; *r* = 0.929 vs FS7). Agreement with FS6 was lower (*r* = 0.842), reflecting systematic differences in boundary definition rather than random error. The hybrid masking approach showed higher regional consistency and lower variability in T1w/T2w-derived myelin estimates (CV = 16.50% vs 40.56% in FastSurfer), with more spatially consistent thickness–myelin interaction patterns in artifact-prone ventral regions. Integrating FastSurfer into the HCP pipeline with hybrid T2w- and T1w-based masking enables stable cortical reconstruction by constraining signal-driven instability while preserving algorithm-dependent spatial organization. These findings highlight the importance of masking as a constraint on instability in cortical reconstruction and support the applicability of HCP-style analysis to clinical and legacy MRI datasets.

## 1. Introduction

In clinical neuroimaging, achieving high-precision cortical reconstruction remains challenging due to practical constraints such as limited scan time, heterogeneous hardware environments, and reliance on standard-resolution (1 mm³) MRI acquisitions. Consequently, there is a growing need for robust methodologies capable of extracting reliable anatomical information from so-called “legacy data”—datasets acquired under routine clinical conditions rather than optimized research protocols (Jack et al., 2008; Marcus et al., 2013). A key challenge is to preserve the anatomical fidelity and reproducibility achieved in large-scale research initiatives, such as the Human Connectome Project (HCP) (Glasser et al., 2013; Miller et al., 2016), while adapting to the reduced signal-to-noise ratio and contrast variability inherent in clinical imaging.

The HCP minimal preprocessing pipeline (hereafter, HCP pipeline), built upon FreeSurfer (Fischl, 2012), represents a widely adopted standard for cortical reconstruction. However, it is primarily optimized for high-resolution acquisitions (e.g., 0.7 mm isotropic), and its behavior under standard-resolution conditions remains less systematically characterized. In particular, subtle resolution- and contrast-dependent effects may emerge in regions susceptible to imaging artifacts, potentially leading to localized inconsistencies in cortical measurements. These effects are not indicative of intrinsic limitations of the HCP framework itself, but rather reflect the interaction between acquisition conditions and preprocessing assumptions.

To address both computational and methodological constraints, recent advances in deep learning-based neuroimaging pipelines have enabled rapid and automated brain segmentation (Roy et al., 2019). Among these approaches, FastSurfer provides a practical solution by combining high computational efficiency with compatibility to FreeSurfer-derived cortical reconstruction, including anatomical segmentation, surface reconstruction, and cortical parcellation (Henschel et al., 2020). In this study, we integrated FastSurfer into the HCP framework to achieve stable and scalable cortical reconstruction under standard-resolution conditions, enabling high-throughput processing on standard CPU-based systems without requiring specialized GPU infrastructure while maintaining compatibility with established FreeSurfer-based workflows.

Despite these advantages, a critical challenge persists in regions affected by susceptibility-induced signal loss, particularly in the orbitofrontal cortex (OFC) and temporal poles (Deichmann et al., 2003). These artifacts are more pronounced in T2-weighted (T2w) images than in T1-weighted (T1w) images and arise from magnetic field inhomogeneities near air–tissue interfaces (Glasser & Van Essen, 2011). Because the HCP pipeline leverages T2w images to refine pial surface placement, such signal dropouts can lead to localized inaccuracies in cortical reconstruction, including spurious surface erosion and distorted morphometric estimates.

In this study, we propose a hybrid cortical reconstruction framework that combines FastSurfer integration with a novel T2w- and T1w-based masking strategy (t2log-hybrid). Our approach begins with automated brain masking using SynthStrip, followed by log-domain refinement of anatomical boundaries. Crucially, we introduce a spatially constrained switching mechanism anchored to the anterior commissure that selectively replaces artifact-prone T2w regions with T1w-derived anatomical information in the orbitofrontal cortex (OFC). This strategy preserves the anatomical advantages of T2w-guided refinement while mitigating its susceptibility to signal dropout.

We hypothesize that such targeted handling of region-specific artifacts can stabilize cortical reconstruction without altering large-scale structural relationships. By systematically comparing pipelines based on FreeSurfer v6.0.1 (FS6), FreeSurfer v7.4.1 (FS7), and FastSurfer, we aim to characterize both global consistency and region-specific deviations. Through complementary analyses of geometric fidelity, regional correlation structure, and vertex-wise coupling patterns, we further investigate how preprocessing differences propagate into derived measures such as thickness–myelin relationships, with particular emphasis on orbitofrontal regions susceptible to signal dropout.

## 2. Materials and methods

### 2.1. Participants and Ethical Approval

This study was conducted in accordance with the Declaration of Helsinki and approved by the Ethics Committee of the Oita University Faculty of Medicine (Approval No. 2599). The analysis used previously acquired MRI data from healthy controls and patients with major depressive disorder. Participants in the original study had provided written informed consent prior to participation. For the present retrospective analysis, information about the study was made publicly available on the department website, and participants were given the opportunity to decline the secondary use of their previously acquired data through an opt-out process. Inclusion required the availability of both T1w and T2w MRI scans. Participants with structural brain abnormalities or neurological comorbidities were excluded. The final dataset included all available participants (n = 10) who met these criteria during the study period, comprising 5 healthy controls and 5 patients with major depressive disorder (5 males and 5 females; age 21–40 years, mean ± SD: 27.8 ± 6.46).

The sample was not selected to test diagnostic group differences, but to evaluate pipeline-dependent effects in standard-resolution structural MRI data. Healthy controls and patients with major depressive disorder were included because these data were available from a previous study and provided standard-resolution T1w and T2w structural MRI acquired under comparable imaging conditions. All primary analyses were therefore performed as within-subject comparisons across reconstruction pipelines, and diagnostic group effects were not modeled.

### 2.2. MRI Acquisition

MRI data were acquired on a 3.0-T system (MAGNETOM Verio, Siemens) using a 32-channel head coil. T1w images were obtained using an MPRAGE sequence (TR = 1900 ms, TE = 2.52 ms, TI = 900 ms, flip angle = 9°, 1.0 mm isotropic resolution). T2w images were acquired using a 3D SPACE sequence (1.0 mm isotropic resolution, GRAPPA factor = 2).

### 2.3. Processing Pipelines and Integration Strategy

All analyses were performed within the HCP minimal preprocessing framework (v5.0.0), incorporating FSL (v6.0.7), FreeSurfer (v6.0.1), and Connectome Workbench (v2.1.0). This configuration is referred to as FS6 and served as the baseline. Two additional configurations were evaluated: FS7 (FreeSurfer v7.4.1 within the HCP framework) and FastSurfer (FastSurfer v2.4.2 with a FreeSurfer v7.4.1 backend). In addition, we developed a hybrid pipeline (LogHybrid), defined as a FastSurfer-based HCP workflow incorporating the t2log-hybrid masking strategy. Segmentation outputs were mapped to HCP-compatible formats, and coordinate consistency was ensured using identity transforms and C-RAS–based alignment. Surface reconstruction was performed in native space to maintain compatibility with the HCP CIFTI workflow.

### 2.4. Hybrid T2w- and T1w-based Masking (t2log-hybrid)

Within the HCP framework, this masking step is applied between the PreFreeSurferPipeline and FreeSurferPipeline stages.

To address susceptibility-induced signal loss in orbitofrontal and anterior temporal regions, we implemented a hybrid anatomical masking strategy. The hybrid masking strategy was primarily designed to stabilize cortical reconstruction at the whole-brain level, whereas the spatial switching mechanism was subsequently introduced to address region-specific signal instability observed in areas such as the orbitofrontal cortex. Initial masks were generated using SynthStrip (Hoopes et al., 2022), followed by refinement using log-domain transformation for T2w images and squared-intensity transformation for T1w images. Although SynthStrip provides robust initial brain extraction, small residual non-brain components and intensity-driven artifacts can persist, particularly low-intensity islands and high-intensity cerebrospinal fluid (CSF) signals adjacent to the cortical surface in T2w images. To address these limitations, these transformations were applied to reduce intensity distribution skew and suppress outlier signals, thereby stabilizing threshold-based separation between brain and non-brain components. Finally, simple morphological operations (erosion followed by dilation) were applied to remove residual isolated components and ensure spatial continuity of the mask. Thresholds were determined using histogram-based criteria (95% confidence interval).

A spatial switching mechanism anchored to the anterior commissure selectively replaced artifact-prone T2w regions with T1w-derived masks in inferior-anterior brain regions. Morphological operations ensured spatial continuity, and the resulting mask was provided to the reconstruction pipeline as an external constraint on pial surface placement during surface reconstruction. For mechanistic validation, a T2w-only masking approach (t2log-strip), representing a simplified version of the hybrid strategy without T1w substitution, was used. The corresponding reconstruction pipeline is referred to as LogStrip. Importantly, these masking strategies operate as input constraints applied prior to surface reconstruction, rather than modifying the reconstruction algorithm itself.

### 2.5. Evaluation Metrics and Statistical Analysis

#### 2.5.1. Global and Regional Consistency

Inter-pipeline consistency was evaluated using Pearson correlation coefficients (*r*) at both vertex-wise and region-wise levels using the Desikan–Killiany–Tourville (DKT) atlas. Global agreement was assessed using pooled vertex distributions, while regional consistency was quantified as the mean of ROI-wise correlations. Correlation-based analyses were performed relative to LogHybrid to facilitate direct assessment of the proposed method, whereas geometric comparisons were conducted relative to FS6 as a conventional baseline.

#### 2.5.2. Geometric Fidelity and Overlap

Geometric accuracy was assessed using surface-to-surface distance (SSD) for white and pial surfaces. Segmentation overlap was quantified using Dice coefficients.

#### 2.5.3. Myelin Signal Stability

Variability in T1w/T2w-derived myelin maps was assessed using the coefficient of variation (CV), calculated within each subject and hemisphere as SD / mean × 100 and then averaged across subjects and hemispheres for each pipeline. This analysis focused on the suppression of non-brain signals such as dura and venous sinuses.

#### 2.5.4. Thickness–myelin association analysis

The relationship between cortical thickness and T1w/T2w-derived myelin estimates was examined using Pearson correlation coefficients as a descriptive measure of association. At the global level, pooled ROI values across all subjects were used to visualize the overall relationship between cortical thickness and myelin estimates within each pipeline. At the regional level, correlation coefficients were computed separately for each ROI across subjects, and their distributions were compared across pipelines. ROIs with zero or invalid values, such as thickness = 0 and myelin = 0, were excluded from ROI-wise correlation analyses. These analyses were intended to characterize the consistency and directionality of thickness–myelin associations rather than to provide formal subject-level inferential statistics.

#### 2.5.5. Vertex-wise interaction analysis

To assess spatially consistent differences between pipelines, an interaction metric was defined at each cortical vertex as the product of cortical thickness and T1w/T2w-derived myelin estimate (thickness × myelin). This metric was used as a descriptive proxy to characterize joint variation patterns rather than as a formal inferential endpoint. For each subject and processing condition, vertex-wise interaction maps were computed on the fsaverage_LR32k surface. Paired comparisons between pipelines were then performed at each vertex across subjects using paired t-statistics to generate spatial maps of within-subject differences, rather than for formal inferential testing. In addition to mean-difference maps, effect sizes were quantified using Cohen’s dz for paired data, calculated as the mean within-subject difference divided by the standard deviation of the within-subject difference at each vertex. These analyses emphasized spatial extent and consistency of effects rather than formal threshold-based statistical inference.

#### 2.5.6. Evaluation of Region-Specific Differences

Region-specific discrepancies were examined in anatomically challenging areas, particularly the orbitofrontal cortex (OFC), cingulate cortex, and paracentral lobule. Reduced correlation with baseline pipelines, especially FS6, was interpreted as reflecting region-specific differences in cortical boundary definition rather than simply indicating poorer reconstruction reliability. Qualitative inspection of cortical surfaces was performed using Freeview and Connectome Workbench, as appropriate, to verify suppression of artifacts without introducing topological distortions. Quantitative analyses and visualization source maps were generated using Python (v3.10) with NumPy, Pandas, SciPy, and NiBabel. Figure rendering was performed using Matplotlib, Surfplot, and Pillow as appropriate.

## 3. Results

### 3.1. Global cortical patterns

Figure 1 shows group-average cortical thickness and T1w/T2w-derived myelin maps across pipelines. Cortical thickness maps exhibited consistent large-scale anatomical patterns, including relatively thin cortex in primary sensorimotor regions and thicker cortex in association areas. However, systematic differences in absolute thickness values were observed. FS6 showed globally higher thickness values, whereas LogHybrid, FastSurfer, and FS7 exhibited lower values, reflecting differences in boundary definition (Table 1). In myelin maps, regional discrepancies were evident. FS7 and FastSurfer showed focal high-intensity signals in medial occipital regions (e.g., calcarine sulcus), whereas LogHybrid showed reduced prominence of these signals. Additionally, FS6 exhibited reduced spatial contrast in regions such as the postcentral gyrus, consistent with differences in pial surface placement. Qualitative inspection of representative cases further illustrated these differences, particularly in regions susceptible to signal-related instability, such as paracentral, orbitofrontal, and medial occipital regions (Supplementary Figure S1). In these examples, FastSurfer, FS7, and FS6 showed pial surface extension beyond the cortical boundary in some locations, whereas LogHybrid remained within the cortical boundary.

**Figure 1.**
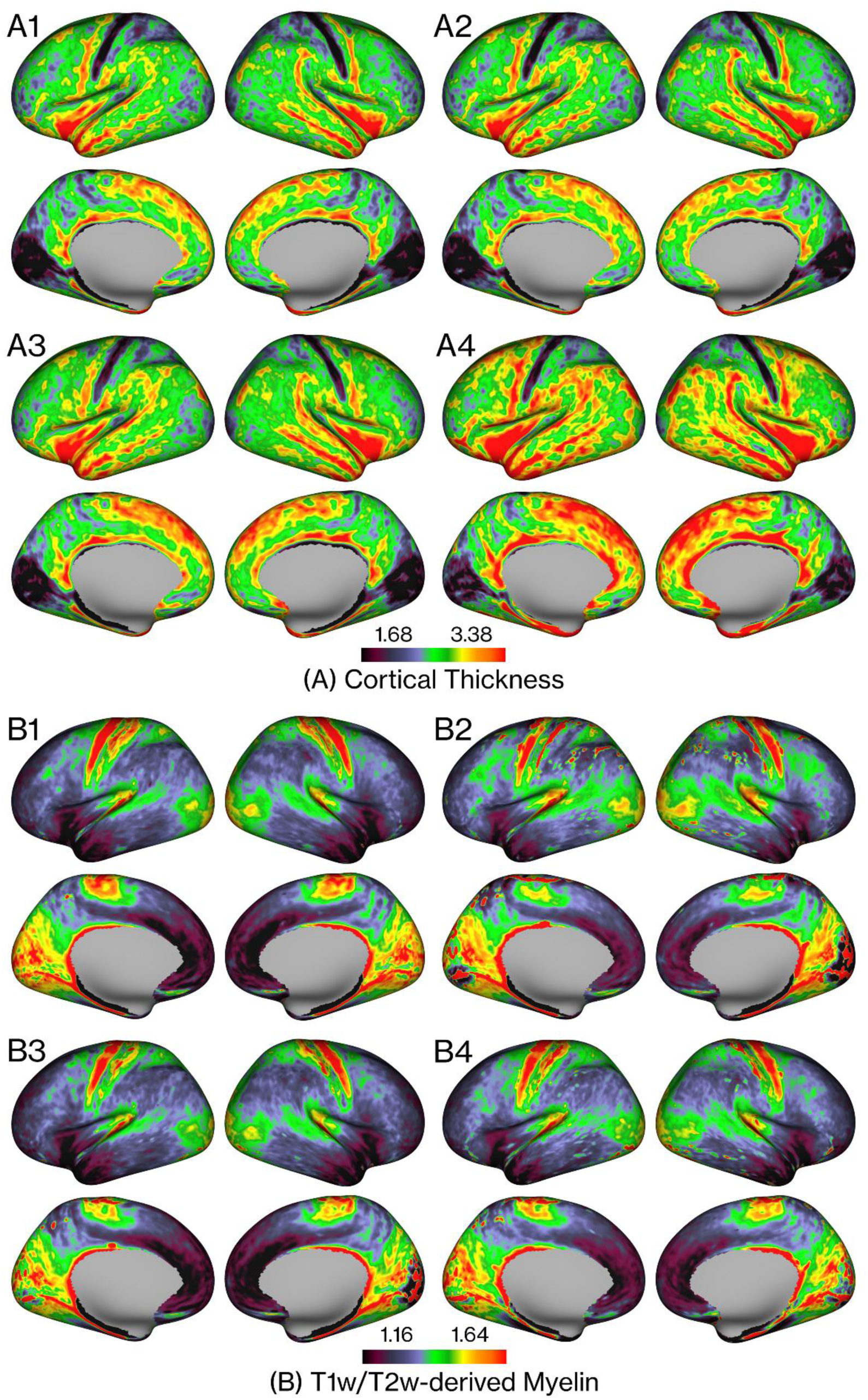
Group-average cortical thickness and myelin maps across HCP-integrated pipelines. (A) Cortical thickness and (B) T1w/T2w-derived myelin maps are shown for LogHybrid, FastSurfer, FS7, and FS6. All pipelines preserve consistent large-scale cortical patterns, but systematic differences in absolute thickness are observed, with FS6 showing higher values. In myelin maps, FS7 and FastSurfer exhibit focal high-intensity signals in medial occipital regions, whereas LogHybrid shows reduced prominence of these signals. FS6 also shows reduced spatial contrast in some regions, consistent with thickness overestimation.

**Table 1.**
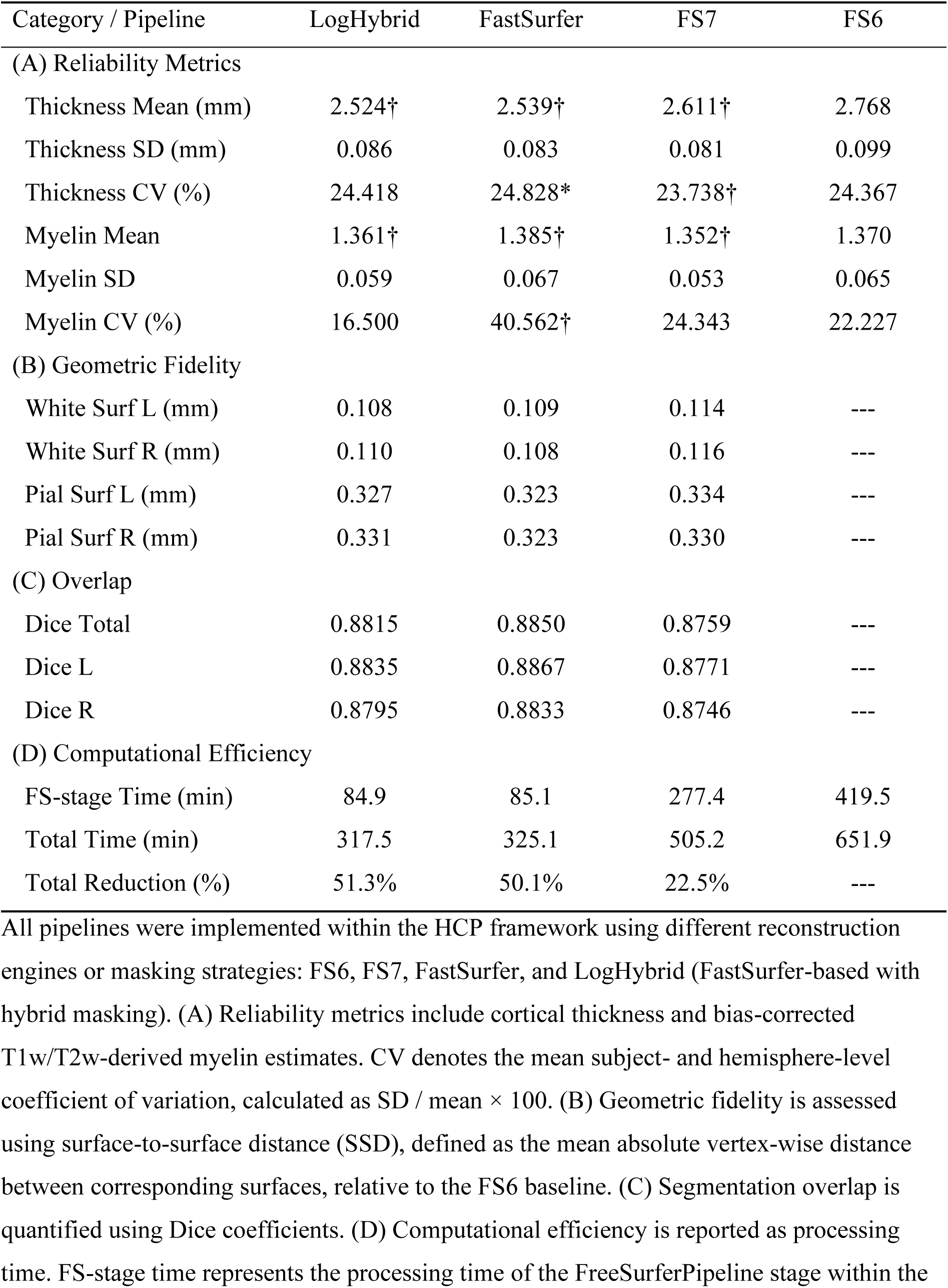
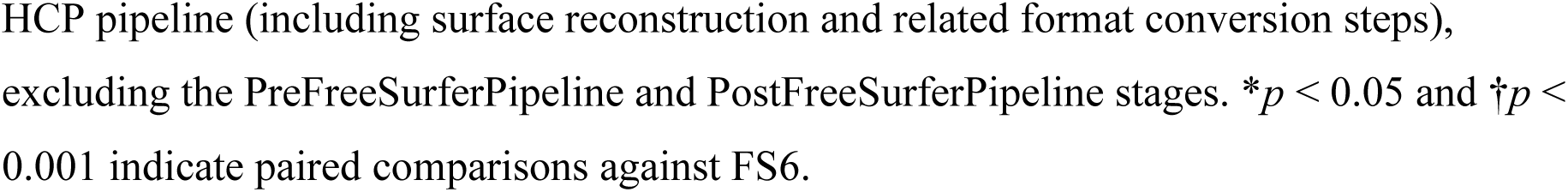
Quantitative performance comparison across cortical reconstruction pipelines.

| Category / Pipeline | LogHybrid | FastSurfer | FS7 | FS6 |
| --- | --- | --- | --- | --- |
| (A) Reliability Metrics |  |  |  |  |
| Thickness Mean (mm) | 2.524† | 2.539† | 2.611† | 2.768 |
| Thickness SD (mm) | 0.086 | 0.083 | 0.081 | 0.099 |
| Thickness CV (%) | 24.418 | 24.828* | 23.738† | 24.367 |
| Myelin Mean | 1.361† | 1.385† | 1.352† | 1.370 |
| Myelin SD | 0.059 | 0.067 | 0.053 | 0.065 |
| Myelin CV (%) | 16.500 | 40.562† | 24.343 | 22.227 |
| (B) Geometric Fidelity |  |  |  |  |
| White Surf L (mm) | 0.108 | 0.109 | 0.114 | --- |
| White Surf R (mm) | 0.110 | 0.108 | 0.116 | --- |
| Pial Surf L (mm) | 0.327 | 0.323 | 0.334 | --- |
| Pial Surf R (mm) | 0.331 | 0.323 | 0.330 | --- |
| (C) Overlap |  |  |  |  |
| Dice Total | 0.8815 | 0.8850 | 0.8759 | --- |
| Dice L | 0.8835 | 0.8867 | 0.8771 | --- |
| Dice R | 0.8795 | 0.8833 | 0.8746 | --- |
| (D) Computational Efficiency |  |  |  |  |
| FS-stage Time (min) | 84.9 | 85.1 | 277.4 | 419.5 |
| Total Time (min) | 317.5 | 325.1 | 505.2 | 651.9 |
| Total Reduction (%) | 51.3% | 50.1% | 22.5% | --- |

All pipelines were implemented within the HCP framework using different reconstruction engines or masking strategies: FS6, FS7, FastSurfer, and LogHybrid (FastSurfer-based with hybrid masking). (A) Reliability metrics include cortical thickness and bias-corrected T1w/T2w-derived myelin estimates. CV denotes the mean subject- and hemisphere-level coefficient of variation, calculated as $SD / \text{mean} \times 100$ . (B) Geometric fidelity is assessed using surface-to-surface distance (SSD), defined as the mean absolute vertex-wise distance between corresponding surfaces, relative to the FS6 baseline. (C) Segmentation overlap is quantified using Dice coefficients. (D) Computational efficiency is reported as processing time. FS-stage time represents the processing time of the FreeSurferPipeline stage within the
HCP pipeline (including surface reconstruction and related format conversion steps), excluding the PreFreeSurferPipeline and PostFreeSurferPipeline stages. \* $p < 0.05$ and † $p < 0.001$ indicate paired comparisons against FS6.

### 3.2. Quantitative performance comparison

Quantitative performance metrics across pipelines are summarized in Table 1. Cortical thickness estimates differed systematically across methods. FS6 showed the highest mean thickness, whereas LogHybrid, FastSurfer, and FS7 exhibited lower values, consistent with differences in boundary definition. Thickness variability (CV) was comparable across methods, indicating similar stability of thickness estimation.

In contrast, substantial differences were observed in myelin signal variability. LogHybrid showed the lowest coefficient of variation (CV = 16.50%), indicating lower signal variability, whereas FastSurfer exhibited markedly higher variability (CV = 40.56%). Geometric fidelity analysis demonstrated that all alternative pipelines maintained close agreement with FS6, with submillimeter surface differences. Overlap analysis showed high Dice coefficients across methods, with only small differences among LogHybrid, FastSurfer, and FS7. In terms of computational efficiency, LogHybrid and FastSurfer reduced total processing time by approximately 50% compared to FS6, while FS7 achieved more modest reductions. These results indicate reduced processing time while maintaining geometric consistency and signal stability.

### 3.3. Vertex-wise agreement and bias analysis

Figure 2A shows hexbin plots of vertex-wise cortical thickness correlations between LogHybrid and the other pipelines, visualizing the density distribution of a large number of cortical vertices. Across all comparisons, the majority of vertices were densely concentrated along the identity line, indicating overall structural agreement. LogHybrid demonstrated very high correlation with FastSurfer (*r* = 0.970), indicating that the geometric properties of the FastSurfer reconstruction are largely preserved after integration of the hybrid masking strategy. Similarly, high agreement was observed with FS7 (*r* = 0.929), consistent with the shared algorithmic foundation between FastSurfer and recent FreeSurfer implementations. In contrast, correlation with FS6 was lower (*r* = 0.842), reflecting a systematic shift in boundary definition rather than random variability. This pattern reflects systematic differences in cortical thickness estimation, with FS6 exhibiting higher cortical thickness estimates in the present analysis. Although the overall correlation structure remained high (*r* > 0.8 across all comparisons), deviations from the identity line were more prominent in specific vertex populations, suggesting region-dependent differences in boundary definition, particularly in anatomically challenging regions such as the orbitofrontal cortex and deep sulci.

**Figure 2.**
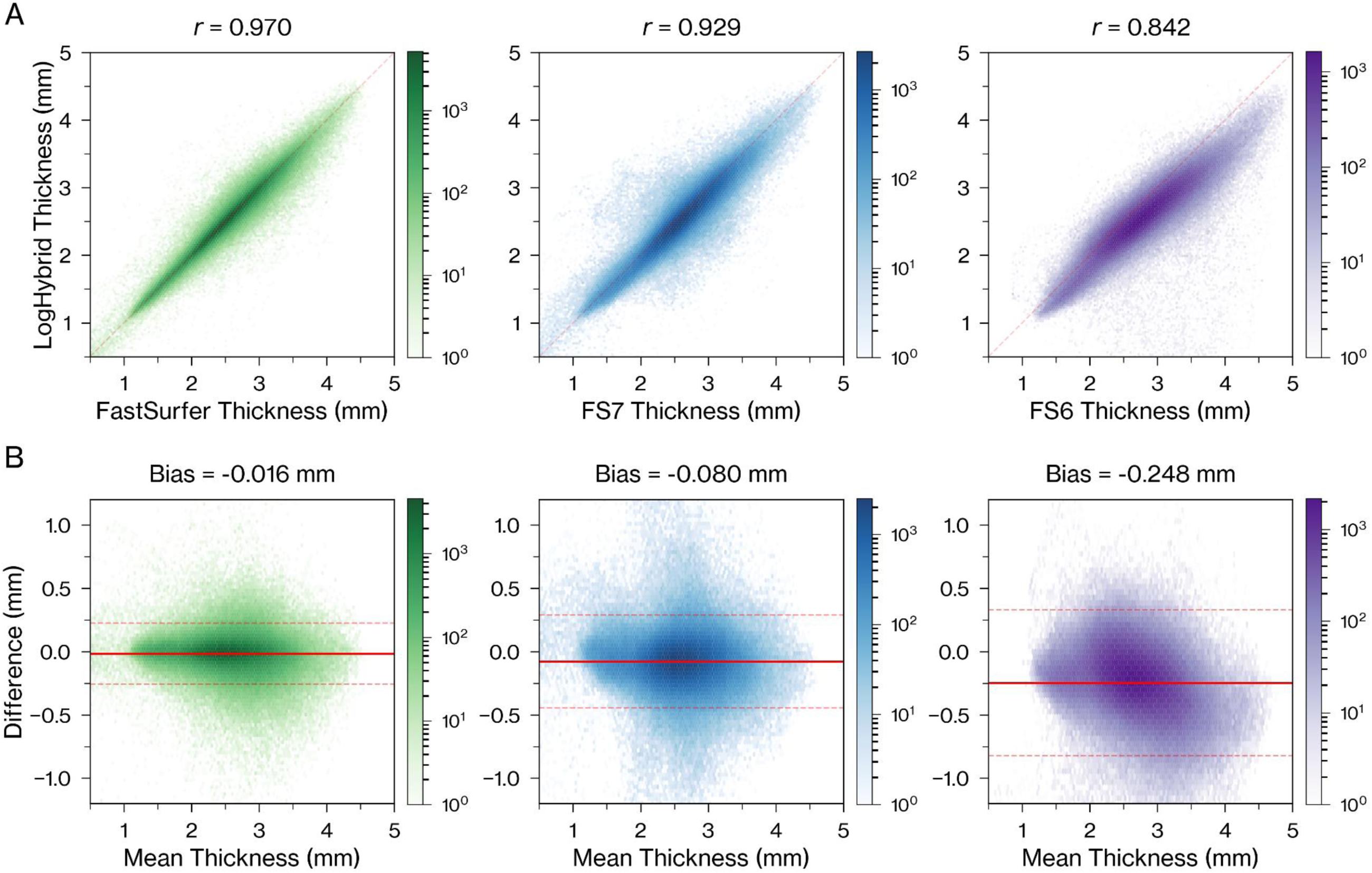
Vertex-wise agreement and bias in cortical thickness across processing pipelines. (A) Hexbin plots show the correspondence of cortical thickness between LogHybrid and each reference pipeline (FastSurfer, FS7, and FS6), computed across all cortical vertices. Agreement is highest for FastSurfer and FS7, whereas correspondence with FS6 is visibly reduced. The dashed red line indicates the line of identity. Hexbin density is shown on a logarithmic scale. (B) Bland–Altman plots show the difference in cortical thickness as a function of the mean thickness for the same comparisons. LogHybrid exhibits minimal bias relative to FastSurfer, modest bias relative to FS7, and a larger negative bias relative to FS6. Solid red lines indicate the mean difference (bias), and dashed red lines indicate the 95% limits of agreement.

Figure 2B presents Bland–Altman plots illustrating systematic bias between pipelines. LogHybrid showed minimal bias relative to FastSurfer (≈ −0.02 mm), indicating near-equivalent thickness estimates. A moderate negative bias was observed relative to FS7 (≈ −0.08 mm), while a pronounced negative bias was observed relative to FS6 (≈ −0.25 mm), reflecting systematic overestimation of cortical thickness in FS6. Notably, this bias was consistently observed across the full range of cortical thickness values rather than being restricted to a specific range, indicating a global shift in boundary definition rather than localized noise effects. The dense concentration of data points and relatively narrow limits of agreement further demonstrate the geometric stability of the proposed method under standard-resolution conditions.

### 3.4. Region-wise consistency analysis

Region-wise consistency relative to LogHybrid is summarized in Table 2. At the global level, all pipelines showed high overall correlation (r > 0.99), indicating strong agreement in large-scale cortical structure. However, differences became more pronounced at the regional level. The mean of ROI-wise correlations was highest for FastSurfer (r = 0.954), indicating that LogHybrid shows regional patterns closely aligned with FastSurfer. In contrast, lower values were observed for FS7 (r = 0.825) and FS6 (r = 0.737), suggesting increasing divergence in regional thickness patterns. This discrepancy indicates that differences between pipelines become apparent at the ROI level rather than in pooled measures. Thus, regional analysis provides a more sensitive assessment of structural consistency across pipelines.

**Table 2.** Region-wise consistency analysis relative to LogHybrid.

| Category / Analysis | FastSurfer | FS7 | FS6 |
| --- | --- | --- | --- |
| Overall Correlation | 0.999 | 0.995 | 0.991 |
| Mean of ROI Correlations | 0.954 | 0.825 | 0.737 |

LogHybrid vs FastSurfer
| Rank | Top 5 Regions ($r$) | Bottom 5 Regions ($r$) |
| --- | --- | --- |
| Rank 1 | R_middletemporal (0.994) | L_lateralorbitofrontal (0.723) |
| Rank 2 | R_inferiortemporal (0.993) | R_caudalanteriorcingulate (0.772) |
| Rank 3 | R_lingual (0.992) | R_medialorbitofrontal (0.785) |
| Rank 4 | L_superiorfrontal (0.992) | L_medialorbitofrontal (0.790) |
| Rank 5 | R_superiorparietal (0.990) | R_paracentral (0.857) |

LogHybrid vs FS7
| Rank | Top 5 Regions ($r$) | Bottom 5 Regions ($r$) |
| --- | --- | --- |
| Rank 1 | L_superiortemporal (0.978) | R_caudalanteriorcingulate (-0.147) |
| Rank 2 | R_inferiorparietal (0.978) | R_medialorbitofrontal (0.386) |
| Rank 3 | L_middletemporal (0.976) | L_medialorbitofrontal (0.543) |
| Rank 4 | L_entorhinal (0.971) | L_caudalanteriorcingulate (0.557) |
| Rank 5 | R_middletemporal (0.971) | R_supramarginal (0.619) |

LogHybrid vs FS6
| Rank | Top 5 Regions ($r$) | Bottom 5 Regions ($r$) |
| --- | --- | --- |
| Rank 1 | L_entorhinal (0.966) | R_caudalanteriorcingulate (-0.199) |
| Rank 2 | R_pericalcarine (0.960) | L_posteriorcingulate (0.015) |
| Rank 3 | R_postcentral (0.947) | R_medialorbitofrontal (0.030) |
| Rank 4 | R_parahippocampal (0.943) | R_paracentral (0.141) |
| Rank 5 | R_cuneus (0.942) | L_paracentral (0.272) |

All comparisons are performed relative to LogHybrid. Global consistency is summarized by overall vertex-wise correlation and the mean of ROI-wise correlations. ROI-wise correlations were computed for each hemisphere and cortical region defined by the DKT atlas using subject-level mean thickness values. Top and bottom regions represent the five highest and lowest ROI-wise Pearson correlation coefficients ( $r$ ), highlighting spatial variation in agreement across pipelines.

Ranking analysis revealed consistent patterns across methods. Regions with high agreement, such as the pericalcarine cortex and entorhinal regions, were shared across pipelines, indicating stable regional thickness patterns in anatomically well-defined areas. In contrast, regions with low agreement showed convergence across pipelines. Notably, orbitofrontal, cingulate, and paracentral regions consistently exhibited reduced correlation. The discrepancy was particularly pronounced in FS6, where correlations approached near-zero or negative values in cingulate, orbitofrontal, and paracentral regions, indicating substantial divergence in regional thickness patterns. These regions are anatomically challenging due to susceptibility-related artifacts and complex cortical geometry. The observed discrepancies, therefore, reflect region-specific limitations of existing pipelines under standard-resolution conditions. In this context, the higher consistency observed with LogHybrid supports the role of targeted hybrid masking in stabilizing regional estimates in these regions. A complete list of ROI-wise correlations is provided in Supplementary Table S1. Consistent with the main analysis, reduced correlations are predominantly observed in orbitofrontal, cingulate, and paracentral regions, particularly in FS6.

### 3.5. Thickness–myelin association analysis

Figure 3 illustrates the association between cortical thickness and T1w/T2w-derived myelin estimates across pipelines. At the global level (Figure 3A), all methods exhibited a positive association between cortical thickness and myelin estimates, reflecting consistent large-scale trends across pipelines. LogHybrid showed a pattern comparable to FastSurfer and FS7. At the regional level (Figure 3B), ROI-wise correlation distributions differed across pipelines. While global associations were positive, ROI-wise correlations showed a broad distribution spanning both positive and negative values. LogHybrid showed a distribution shifted toward more negative correlations relative to FS6, indicating differences in the spatial expression of the thickness–myelin relationship. These analyses are descriptive and reflect pooled and ROI-wise association patterns rather than formal subject-level inference.

**Figure 3.**
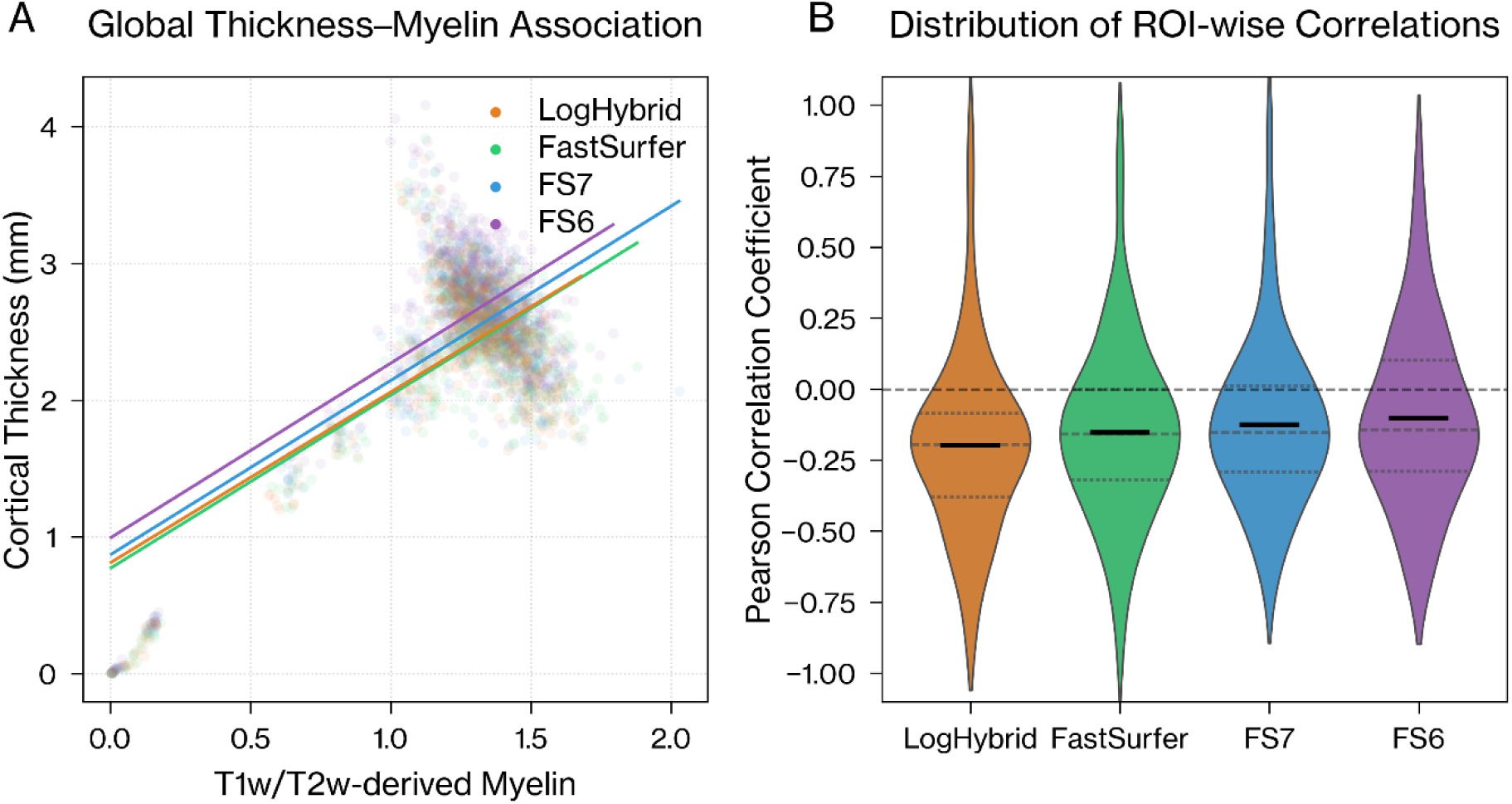
Global and regional thickness–myelin association across processing pipelines. (A) Global thickness–myelin association: pooled scatter plots of cortical thickness and T1w/T2w-derived myelin across all subjects and DKT regions for each pipeline. Solid lines indicate linear regression fits, showing a consistent positive association across all methods. (B) Distribution of ROI-wise correlations: violin plots of Pearson correlation coefficients calculated separately for each hemisphere and DKT region using subject-level mean values. All pipelines were implemented within the HCP framework using different reconstruction engines or masking strategies (LogHybrid, FastSurfer, FS7, and FS6). While global associations are positive, ROI-wise correlations show a distribution shifted toward negative values, reflecting spatially structured inverse relationships between thickness and myelin. Compared with FS6, LogHybrid shows a distribution shifted toward more negative values, indicating a more consistent expression of this regional pattern.

### 3.6. Vertex-wise effect size analysis of the thickness–myelin interaction metric

Figure 4 presents vertex-wise effect size maps (Cohen’s *d_z_*) derived from paired comparisons of the thickness–myelin interaction metric across pipelines. Panels A and B compare LogHybrid and LogStrip with the FS6 baseline, whereas panel C directly compares LogHybrid with LogStrip. Both LogHybrid and LogStrip showed predominantly negative effects relative to FS6 across widespread cortical regions. LogHybrid showed a slightly broader extent of negative effect sizes in orbitofrontal regions, whereas LogStrip showed a more localized pattern.

**Figure 4.**
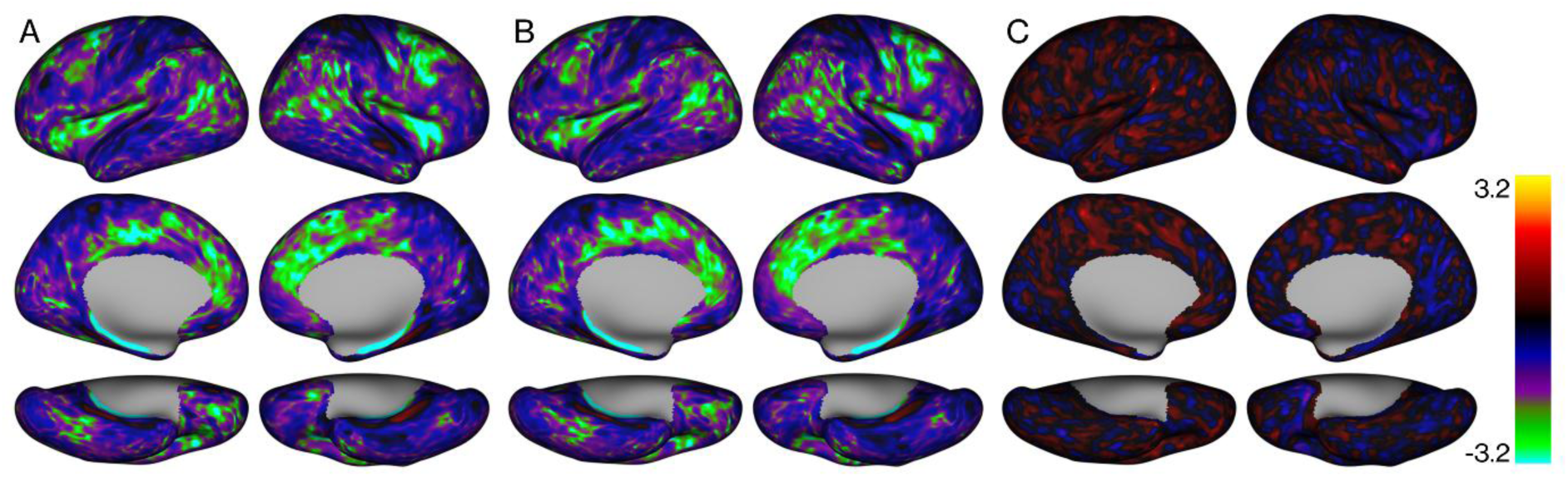
Vertex-wise effect size maps (Cohen’s *d_z_*) of the thickness–myelin interaction metric across pipeline comparisons. (A) LogHybrid vs FS6; (B) LogStrip vs FS6; and (C) LogHybrid vs LogStrip. Panels A and B show effect size maps relative to the FS6 baseline, whereas panel C directly compares the hybrid T2w–T1w masking strategy with the T2w-only masking strategy. Both LogHybrid and LogStrip exhibit predominantly negative effects relative to FS6 across widespread cortical regions. The direct LogHybrid–LogStrip comparison shows more subtle differences, with localized effects in inferior-anterior cortical regions.

The direct LogHybrid–LogStrip comparison (Figure 4C) suggested subtle, region-specific differences primarily in inferior-anterior regions, including the OFC. These differences were less apparent in raw mean-difference maps but clearer in the standardized effect size representation, indicating differences in spatial consistency rather than absolute magnitude. These patterns were consistent with the distribution of the interaction metric (Supplementary Figure S2). Thresholded maps (*p* < 0.001) showed a similar spatial distribution (Supplementary Figure S3).

## 4. Discussion

### 4.1. Overview of Main Findings

This study demonstrates that integrating a hybrid masking approach into the HCP pipeline with FastSurfer preserves global cortical organization while modulating regional differences under standard-resolution conditions. Across all pipelines, large-scale patterns of cortical thickness and myelin were highly consistent, indicating that global structure is preserved. Systematic differences in mean cortical thickness were observed across pipelines, particularly relative to FS6, reflecting differences in the underlying reconstruction algorithms. These differences were not random but reflected consistent shifts in pial surface placement.

In contrast, reduced variability in myelin estimates indicates a distinct effect of the hybrid masking strategy. Specifically, lower CV values suggest suppression of non-brain signal contamination and stabilization of pial surface estimation in artifact-prone regions. In this sense, the observed results reflect two separable effects: algorithm-dependent shifts in mean values and masking-driven reduction of variability. Accordingly, pial surface placement is determined by the reconstruction algorithm, while masking acts to constrain signal-driven instability without altering the global structure.

### 4.2. Global Consistency and Regional Variability

A key finding of this study is the dissociation between global agreement and regional variability across reconstruction pipelines. Although all methods demonstrated high overall correspondence in cortical thickness, as reflected by strong vertex-wise correlations, this apparent agreement masked substantial discrepancies at the regional level. ROI-wise analyses revealed pronounced divergence in specific cortical areas, particularly in orbitofrontal, cingulate, and paracentral regions, where correlations with FS6 were markedly reduced. This pattern indicates that global summary metrics alone may be insufficient to capture meaningful differences in cortical reconstruction, particularly when discrepancies are spatially localized. These findings underscore the importance of disentangling global structural preservation from local reconstruction variability when evaluating neuroimaging pipelines. Notably, such regional differences are consistent with the effects of masking constraints on pial surface estimation, rather than direct alterations of image intensities.

### 4.3. Resolution-Dependent Behavior of FS6

The systematic elevation of cortical thickness observed in FS6 is consistent with the resolution-dependent behavior expected when a pipeline designed for high-resolution HCP-style acquisitions is applied to standard-resolution data. Under standard-resolution conditions, reduced gray–white contrast and increased partial volume effects can compromise boundary delineation, potentially contributing to overestimation of cortical thickness. In particular, susceptibility-prone regions such as the orbitofrontal cortex are especially vulnerable to such effects, as signal dropout and distortion further degrade boundary contrast. These observations are supported by both qualitative and quantitative findings, including reduced regional correlations and visual evidence of surface displacement, indicating that the increased thickness values reflect resolution-dependent limitations of the FS6-based reconstruction framework.

### 4.4. Thickness–Myelin Interaction Patterns

The interaction between cortical thickness and myelin provides a complementary perspective on reconstruction behavior. In the present study, a broadly positive association was observed at the global level across pipelines. This pattern likely reflects methodological factors related to thickness estimation and T1w/T2w-based myelin mapping, rather than a direct biological relationship. However, this global trend concealed substantial variability at the regional level. ROI-wise analyses revealed a wide distribution of correlation values, with certain pipelines exhibiting shifts in the distribution of associations.

Importantly, at the regional level, more negative associations are consistent with previously described interactions between cortical myelin and cortical thickness, in which more heavily myelinated cortex has been described as tending to be thinner after accounting for curvature-related effects (Glasser & Van Essen, 2011). This interpretation is also compatible with multimodal cortical organization, in which highly myelinated primary sensory regions differ systematically from less myelinated association regions (Glasser et al., 2016). In this context, the relative shift toward more negative correlations observed in LogHybrid compared with FS6 may reflect reduced influence of non-brain signal contributions, which can affect myelin estimates and obscure such spatial relationships. Thus, the observed changes likely arise from a combination of algorithm-dependent shifts in mean cortical thickness and masking-driven reductions in signal variability. Within this framework, interaction-based metrics provide a useful means of characterizing such differences beyond global summaries.

### 4.5. Mechanistic Interpretation of Hybrid Masking

Spatially consistent differences were observed across the cortex, which are most plausibly explained by the interaction between susceptibility-related artifacts and pial surface estimation. Regions such as the orbitofrontal cortex (OFC), where susceptibility effects are well known, provide a representative context for interpreting these patterns. The OFC is particularly vulnerable to signal dropout in T2w imaging due to magnetic field inhomogeneities near air–tissue interfaces, leading to reduced contrast and instability in pial surface placement (Deichmann et al., 2003; Glasser & Van Essen, 2011; Fischl, 2012). Because the HCP pipeline uses T2w information to refine the pial surface, such signal degradation can propagate into local inaccuracies in cortical boundary estimation. Although T2w information is used across multiple reconstruction pipelines to refine pial surface placement, masking operates at a different stage by constraining the input signal prior to surface estimation. This, in turn, influences how such refinements are applied.

In this context, both LogHybrid and the T2w-based masking approach (LogStrip) showed consistent reductions in region-specific deviations relative to FS6, indicating that masking plays a key role in limiting the influence of non-brain or unstable signals on pial surface estimation. This suggests that the primary contribution of the masking strategy is not to redefine the reconstruction itself, but to constrain instability arising from signal degradation under standard-resolution conditions.

Notably, spatially consistent effects were not limited to regions directly affected by masking. Similar patterns were also observed across different reconstruction configurations, including FS7 and FastSurfer, indicating that part of the spatial structure arises from the reconstruction algorithm itself. In this context, masking does not generate these patterns de novo, but interacts with an existing algorithm-dependent spatial distribution by constraining instability in regions susceptible to signal degradation. Accordingly, the observed spatial differences reflect the combined influence of reconstruction algorithms and masking constraints, with the former shaping the baseline distribution and the latter modulating its expression.

Taken together, these findings suggest that masking contributes substantially to the observed effects by stabilizing pial surface estimation, while the hybrid strategy provides a more controlled and anatomically constrained implementation of this principle under challenging imaging conditions. Importantly, the spatial switching mechanism anchored to the anterior commissure does not introduce artificial discontinuities or boundary artifacts, as evidenced by the absence of boundary-aligned deviations and the small mean differences observed across the cortex. These observations indicate that the switching mechanism functions as a controlled constraint rather than a source of geometric distortion.

In conventional pipelines, masking is primarily treated as a preprocessing step for brain extraction, whereas in the present approach it is used as an explicit constraint to guide surface reconstruction. Although the present implementation was evaluated within a FastSurfer-integrated framework, the masking strategy itself operates as an input-level constraint applied prior to surface reconstruction. In principle, this approach may be applicable to other reconstruction pipelines, including conventional FreeSurfer-based implementations. However, the extent and spatial characteristics of its effects may vary depending on how T2w information is incorporated within each pipeline. Therefore, further evaluation across different reconstruction frameworks would be required to fully characterize its applicability.

### 4.6. Methodological Implications

Previous methodological studies have shown that preprocessing pipelines, reconstruction algorithms, and software versions can systematically influence cortical surface analyses and morphometric estimates. Bhagwat et al. (2021) demonstrated that the choice of preprocessing pipeline can affect neuroimaging cortical surface analyses, while Kharabian Masouleh et al. (2020) showed that cortical thickness estimates differ across major morphometric pipelines, including FreeSurfer, CIVET, and CAT. Filip et al. (2022) further showed that FreeSurfer version and HCP-specific processing variants can affect downstream statistical outcomes. Together, these studies underscore the importance of using fixed, version-documented workflows rather than treating reconstruction pipelines as interchangeable. The present study follows this pipeline-comparison perspective, but differs by evaluating a FastSurfer-integrated HCP workflow and an input-level hybrid masking constraint under standard-resolution conditions.

An important methodological aspect of this study is the use of an interaction-based metric (thickness × myelin) combined with effect size mapping (Cohen’s *d_z_*) to characterize spatially consistent differences between pipelines. Unlike conventional correlation analyses, which summarize associations either globally or at the ROI level, this approach captures vertex-wise patterns of within-subject differences, emphasizing spatial coherence rather than statistical significance alone. This distinction is particularly relevant in the present context, where preprocessing differences produce subtle but consistent shifts that may not manifest as large mean differences.

By focusing on effect size rather than thresholded statistical maps, the analysis avoids overreliance on arbitrary significance thresholds and instead highlights regions where differences are consistently expressed across subjects. This is particularly important for detecting spatially distributed effects, where the primary distinction between methods lies in pattern coherence rather than magnitude.

Importantly, the interaction metric used here should not be interpreted as a direct measure of biological coupling or reconstruction accuracy, but rather as a proxy reflecting how preprocessing choices influence the joint distribution of cortical thickness and myelin estimates. Within this framework, the observed interaction patterns can be understood as downstream consequences of boundary definition, providing a sensitive readout of how localized biases propagate into higher-level representations of cortical organization. This property also suggests potential utility for quality control applications, although such use was not the primary focus of the present study.

### 4.7. Limitations

Several limitations should be considered when interpreting these findings. First, the small sample size limits the generalizability of the findings and warrants validation in larger cohorts. Because the sample included both healthy controls and patients with major depressive disorder, potential diagnostic differences in the magnitude of masking-related correction cannot be excluded. However, the present study was not designed or powered to assess diagnostic group effects, and all analyses focused on within-subject, pipeline-dependent differences. Because region-specific reconstruction failures may vary across individuals, such effects can be attenuated in large group-average maps; therefore, future validation should combine larger cohorts with subject-level quality-control assessments.

Second, the HCP pipeline was originally optimized for high-resolution acquisitions (e.g., 0.7 mm), and its behavior under standard-resolution conditions is not fully characterized. The present results, therefore, reflect resolution-dependent effects rather than intrinsic limitations of the underlying algorithms, and no direct comparison with high-resolution data was performed.

Third, because T2-weighted images are incorporated after alignment to the T1w space, all processing steps that utilize T2w information inherently depend on the accuracy of T1w–T2w registration. This includes both pial surface refinement in conventional pipelines (e.g., FS6 and FS7) and the present hybrid masking approach, although the impact may differ depending on how T2w information is incorporated.

Fourth, the interaction-based metric used in this study serves as a proxy for joint variation in cortical thickness and myelin and should not be interpreted as a direct measure of biological coupling or reconstruction accuracy.

## 5. Conclusion

This study demonstrates that cortical reconstruction under standard-resolution conditions is stabilized by constraining signal-driven instability through a hybrid masking strategy while preserving algorithm-dependent spatial organization. These findings highlight the role of masking in constraining instability during cortical reconstruction and support the applicability of HCP-style analysis to standard-resolution MRI.

## Funding

This work was supported by JSPS KAKENHI Grant Number 18K15487. MRI data used in this study were acquired as part of this funded project.

## CRediT authorship contribution statement

Koji Hatano: Conceptualization, Methodology, Software, Formal analysis, Investigation, Resources, Data curation, Visualization, Writing – original draft, Writing – review & editing, Project administration, Funding acquisition. Hirofumi Hirakawa: Investigation, Resources, Writing – review & editing. Hiroyuki Matsuda: Investigation, Resources, Writing – review & editing. Nobuhiko Hoaki: Supervision, Writing – review & editing. Takeshi Terao: Supervision, Writing – review & editing. Tsuyoshi Shimomura: Investigation, Resources, Supervision, Writing – review & editing.

## Data and code availability statement

The hybrid masking workflow used in this study is available as t2log-hybrid v4.11 at https://github.com/koji-hatano1/t2log-hybrid. The T2w-based masking variant used as the LogStrip comparison is available as t2log-strip v3.12 at https://github.com/koji-hatano1/t2log-strip. The main analysis scripts and implementation notes used in this study are publicly available at: https://github.com/koji-hatano1/hcp_loghybrid_analysis. Additional implementation details are available from the corresponding author upon reasonable request. The datasets generated and/or analyzed during the current study are not publicly available due to institutional and ethical restrictions but are available from the corresponding author upon reasonable request, subject to approval by the relevant ethics committee.

## Declaration of generative AI and AI-assisted technologies in the manuscript preparation process

During the preparation of this work, the authors used ChatGPT (OpenAI) for language editing and organization. The authors reviewed and edited the content as needed and take full responsibility for the content of the manuscript.

## Declaration of competing interest

The authors declare no competing interests.

## Supplementary Materials

**Supplementary Table S1.**
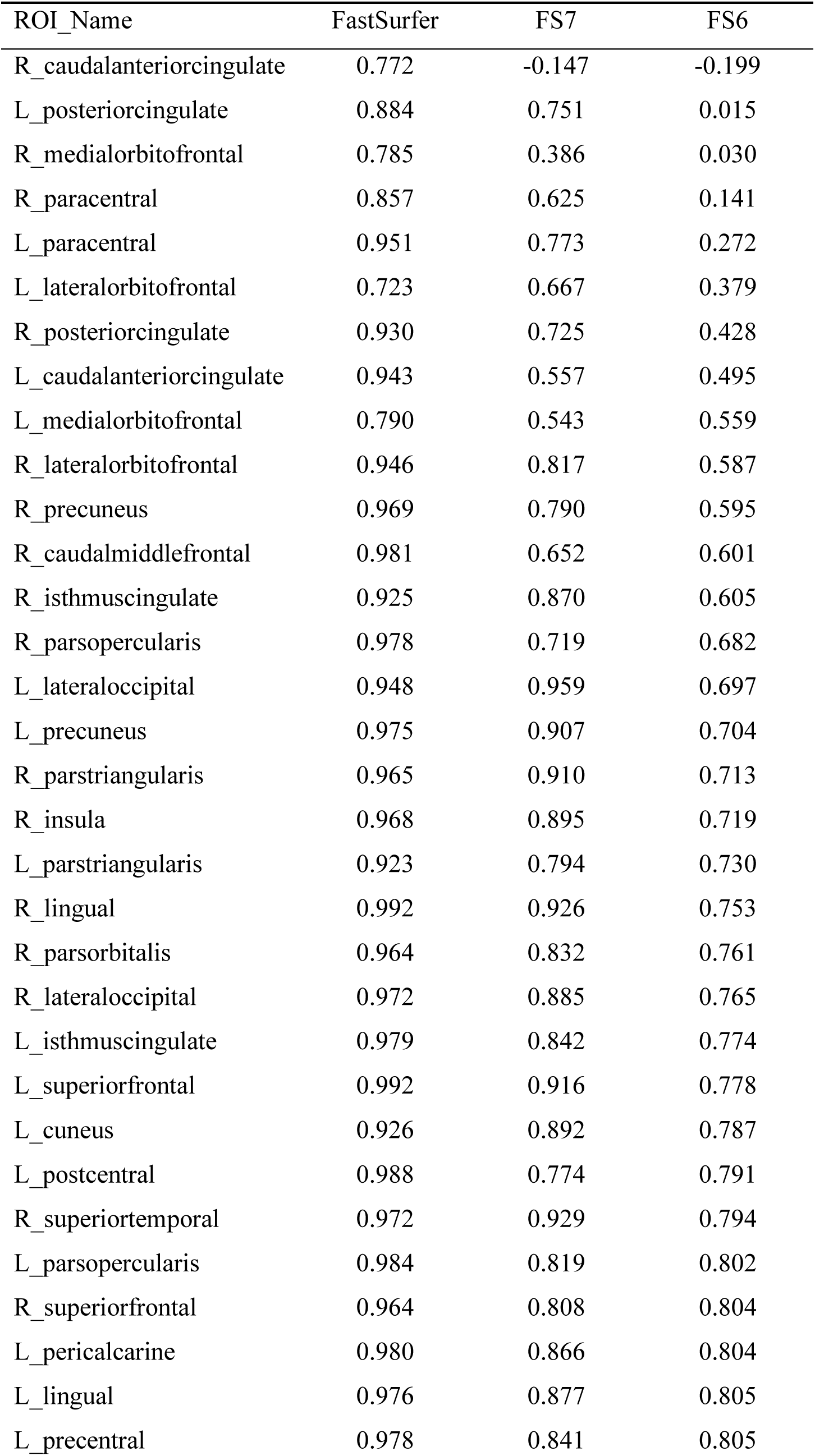
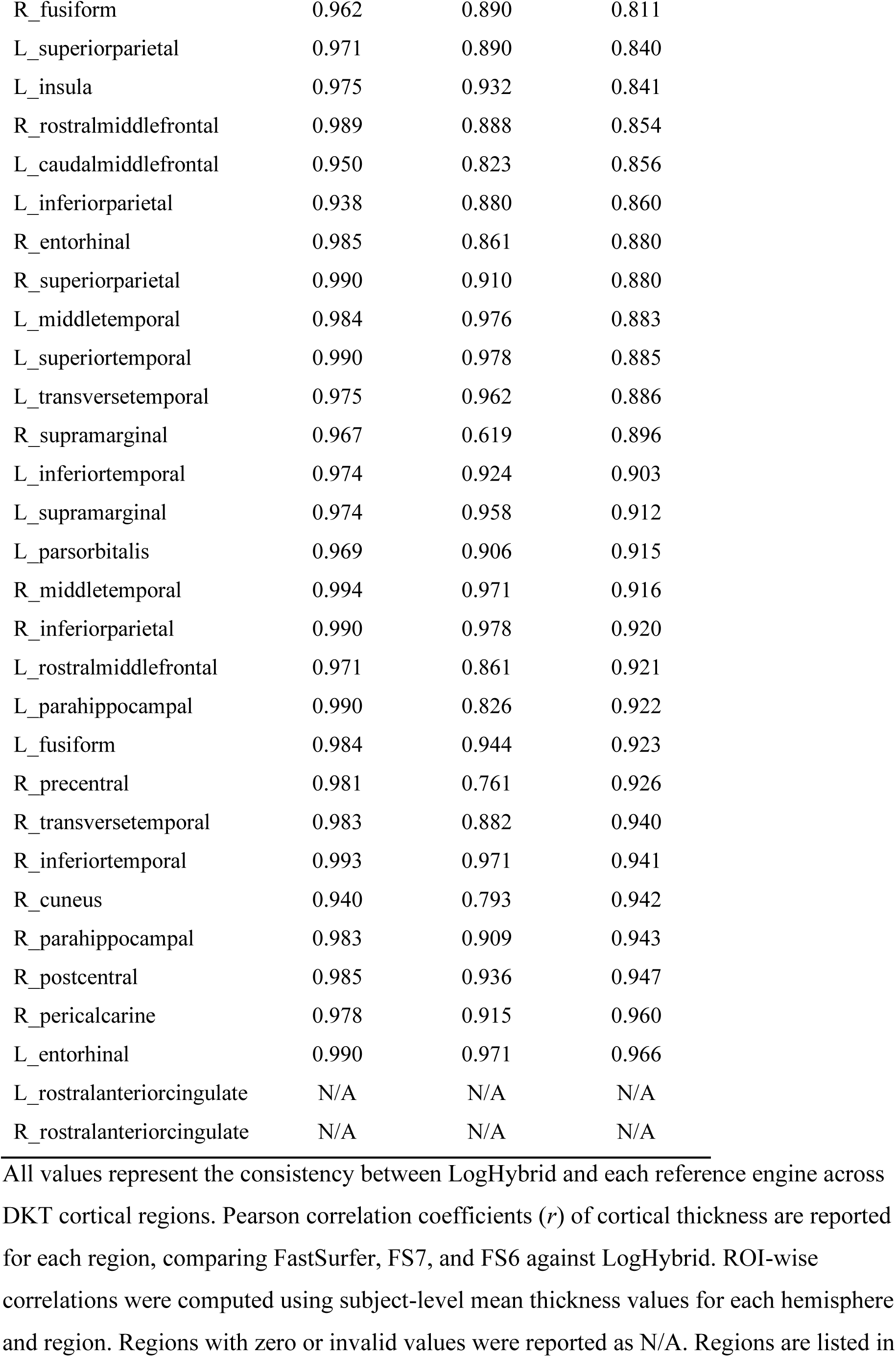
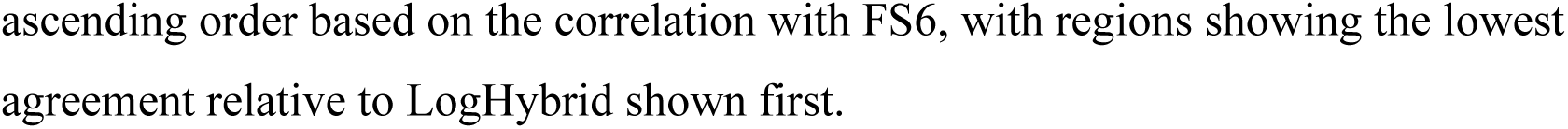
ROI-wise correlation coefficients relative to LogHybrid.

| ROI_Name | FastSurfer | FS7 | FS6 |
| --- | --- | --- | --- |
| R_caudalanteriorcingulate | 0.772 | -0.147 | -0.199 |
| L_posteriorcingulate | 0.884 | 0.751 | 0.015 |
| R_medialorbitofrontal | 0.785 | 0.386 | 0.030 |
| R_paracentral | 0.857 | 0.625 | 0.141 |
| L_paracentral | 0.951 | 0.773 | 0.272 |
| L_lateralorbitofrontal | 0.723 | 0.667 | 0.379 |
| R_posteriorcingulate | 0.930 | 0.725 | 0.428 |
| L_caudalanteriorcingulate | 0.943 | 0.557 | 0.495 |
| L_medialorbitofrontal | 0.790 | 0.543 | 0.559 |
| R_lateralorbitofrontal | 0.946 | 0.817 | 0.587 |
| R_precuneus | 0.969 | 0.790 | 0.595 |
| R_caudalmiddlefrontal | 0.981 | 0.652 | 0.601 |
| R_isthmuscingulate | 0.925 | 0.870 | 0.605 |
| R_parsopercularis | 0.978 | 0.719 | 0.682 |
| L_lateraloccipital | 0.948 | 0.959 | 0.697 |
| L_precuneus | 0.975 | 0.907 | 0.704 |
| R_parstriangularis | 0.965 | 0.910 | 0.713 |
| R_insula | 0.968 | 0.895 | 0.719 |
| L_parstriangularis | 0.923 | 0.794 | 0.730 |
| R_lingual | 0.992 | 0.926 | 0.753 |
| R_parsorbitalis | 0.964 | 0.832 | 0.761 |
| R_lateraloccipital | 0.972 | 0.885 | 0.765 |
| L_isthmuscingulate | 0.979 | 0.842 | 0.774 |
| L_superiorfrontal | 0.992 | 0.916 | 0.778 |
| L_cuneus | 0.926 | 0.892 | 0.787 |
| L_postcentral | 0.988 | 0.774 | 0.791 |
| R_superiortemporal | 0.972 | 0.929 | 0.794 |
| L_parsopercularis | 0.984 | 0.819 | 0.802 |
| R_superiorfrontal | 0.964 | 0.808 | 0.804 |
| L_pericalcarine | 0.980 | 0.866 | 0.804 |
| L_lingual | 0.976 | 0.877 | 0.805 |
| L_precentral | 0.978 | 0.841 | 0.805 |
| R_fusiform | 0.962 | 0.890 | 0.811 |
| L_superiorparietal | 0.971 | 0.890 | 0.840 |
| L_insula | 0.975 | 0.932 | 0.841 |
| R_rostralmiddlefrontal | 0.989 | 0.888 | 0.854 |
| L_caudalmiddlefrontal | 0.950 | 0.823 | 0.856 |
| L_inferiorparietal | 0.938 | 0.880 | 0.860 |
| R_entorhinal | 0.985 | 0.861 | 0.880 |
| R_superiorparietal | 0.990 | 0.910 | 0.880 |
| L_middletemporal | 0.984 | 0.976 | 0.883 |
| L_superiortemporal | 0.990 | 0.978 | 0.885 |
| L_transversetemporal | 0.975 | 0.962 | 0.886 |
| R_supramarginal | 0.967 | 0.619 | 0.896 |
| L_inferiortemporal | 0.974 | 0.924 | 0.903 |
| L_supramarginal | 0.974 | 0.958 | 0.912 |
| L_parsorbitalis | 0.969 | 0.906 | 0.915 |
| R_middletemporal | 0.994 | 0.971 | 0.916 |
| R_inferiorparietal | 0.990 | 0.978 | 0.920 |
| L_rostralmiddlefrontal | 0.971 | 0.861 | 0.921 |
| L_parahippocampal | 0.990 | 0.826 | 0.922 |
| L_fusiform | 0.984 | 0.944 | 0.923 |
| R_precentral | 0.981 | 0.761 | 0.926 |
| R_transversetemporal | 0.983 | 0.882 | 0.940 |
| R_inferiortemporal | 0.993 | 0.971 | 0.941 |
| R_cuneus | 0.940 | 0.793 | 0.942 |
| R_parahippocampal | 0.983 | 0.909 | 0.943 |
| R_postcentral | 0.985 | 0.936 | 0.947 |
| R_pericalcarine | 0.978 | 0.915 | 0.960 |
| L_entorhinal | 0.990 | 0.971 | 0.966 |
| L_rostralanteriorcingulate | N/A | N/A | N/A |
| R_rostralanteriorcingulate | N/A | N/A | N/A |

All values represent the consistency between LogHybrid and each reference engine across DKT cortical regions. Pearson correlation coefficients ( $r$ ) of cortical thickness are reported for each region, comparing FastSurfer, FS7, and FS6 against LogHybrid. ROI-wise correlations were computed using subject-level mean thickness values for each hemisphere and region. Regions with zero or invalid values were reported as N/A. Regions are listed in
ascending order based on the correlation with FS6, with regions showing the lowest agreement relative to LogHybrid shown first.

**Supplementary Figure S1.**
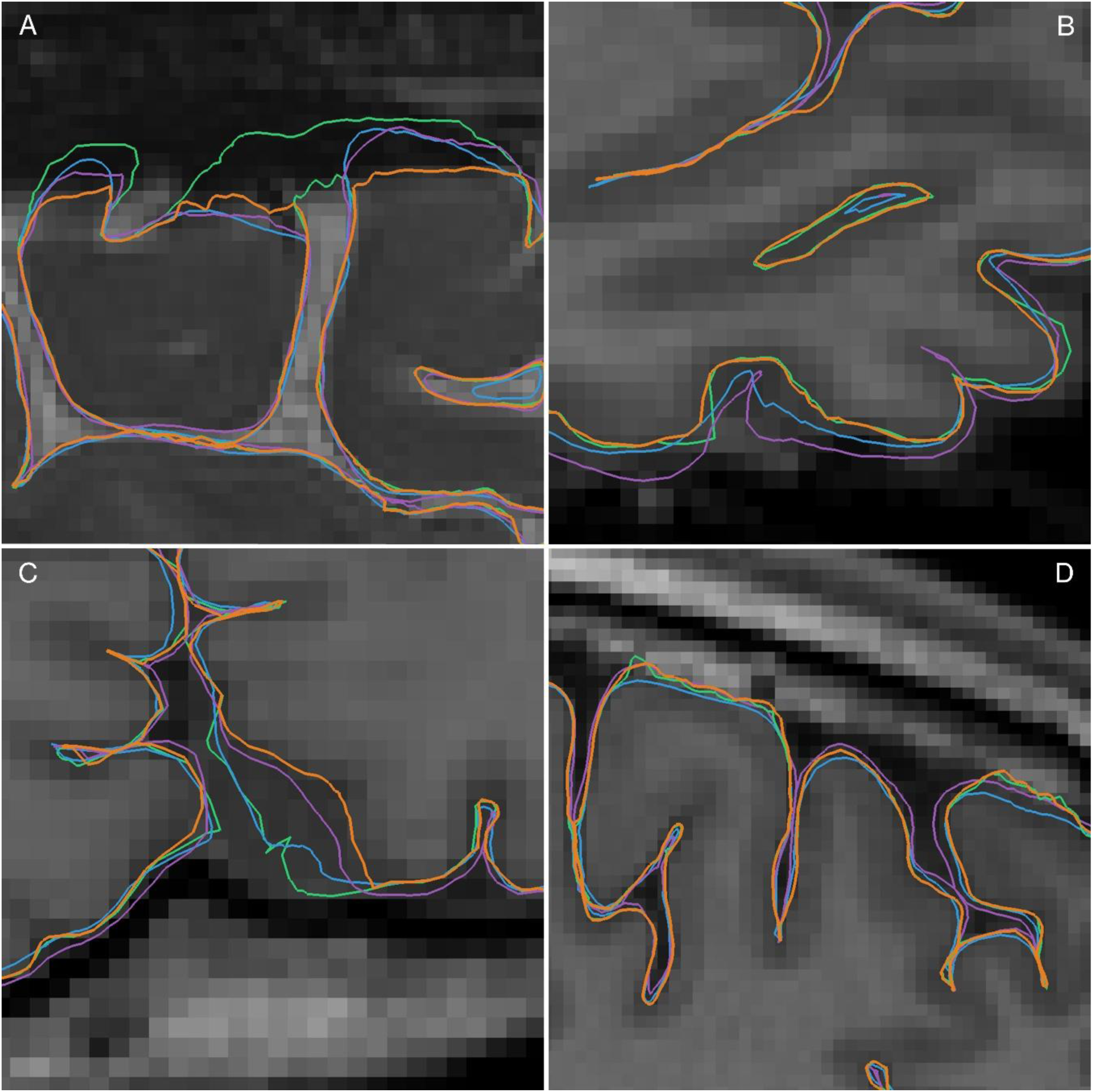
Examples of pial surface placement across reconstruction pipelines. Freeview visualizations of cortical surfaces are shown on T2-weighted (A) and T1-weighted (B–D) images. Orange: LogHybrid, green: FastSurfer, cyan: FS7, purple: FS6. Representative examples are shown in panels A–D. (A) Paracentral region: in FastSurfer, FS7, and FS6, the pial surface extends beyond the cortical boundary, whereas LogHybrid remains within the cortical contour. (B) Orbitofrontal cortex: FS6 shows a systematically thicker boundary relative to other methods, consistent with differences in pial surface definition. (C) Medial occipital region adjacent to venous sinuses: FastSurfer, FS7, and FS6 exhibit surface extension into non-cortical signal regions, while LogHybrid suppresses these effects. (D) Example of apparent boundary truncation: although LogHybrid appears more constrained due to masking, its surface placement remains comparable to FastSurfer, indicating that no artificial boundary displacement is introduced. Across panels, LogHybrid constrains surface placement within the cortical boundary, with reduced influence of non-brain signals observed in regions susceptible to signal-related instability.

**Supplementary Figure S2.**
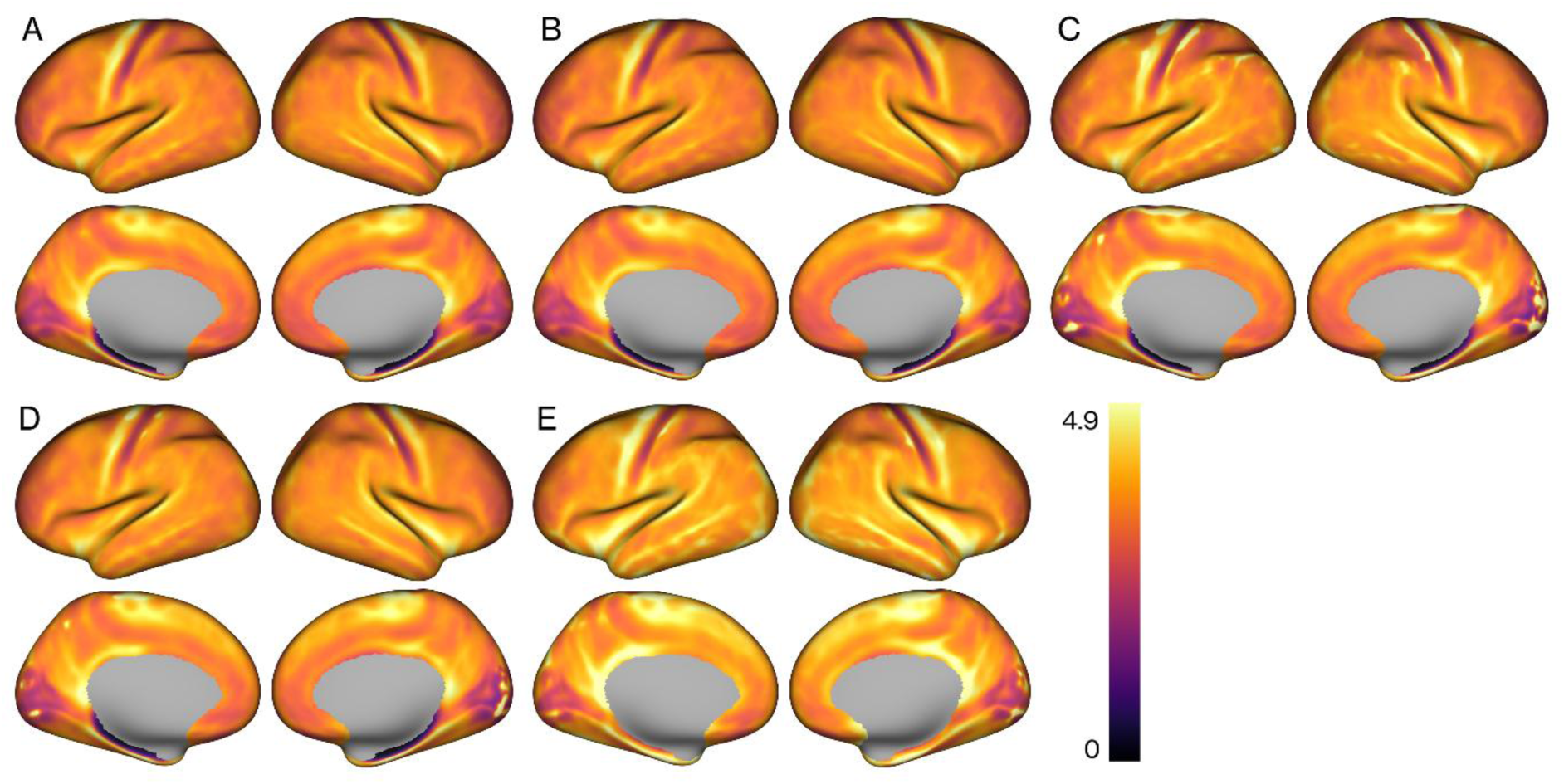
Group-average maps of the thickness–myelin interaction metric across processing pipelines. Mean maps of the interaction metric (cortical thickness × T1w/T2w-derived myelin) are shown using a common color scale for (A) LogHybrid, (B) LogStrip, (C) FastSurfer, (D) FS7, and (E) FS6. FS6 exhibits a globally elevated interaction level, indicating a systematic shift in the joint distribution of cortical thickness and myelin. In contrast, FastSurfer and FS7 show more spatially localized high-intensity patterns that appear as focal, spot-like fluctuations, particularly in anatomically challenging regions. LogHybrid and LogStrip display more spatially coherent and constrained patterns, with reduced global elevation compared to FS6.

**Supplementary Figure S3.**
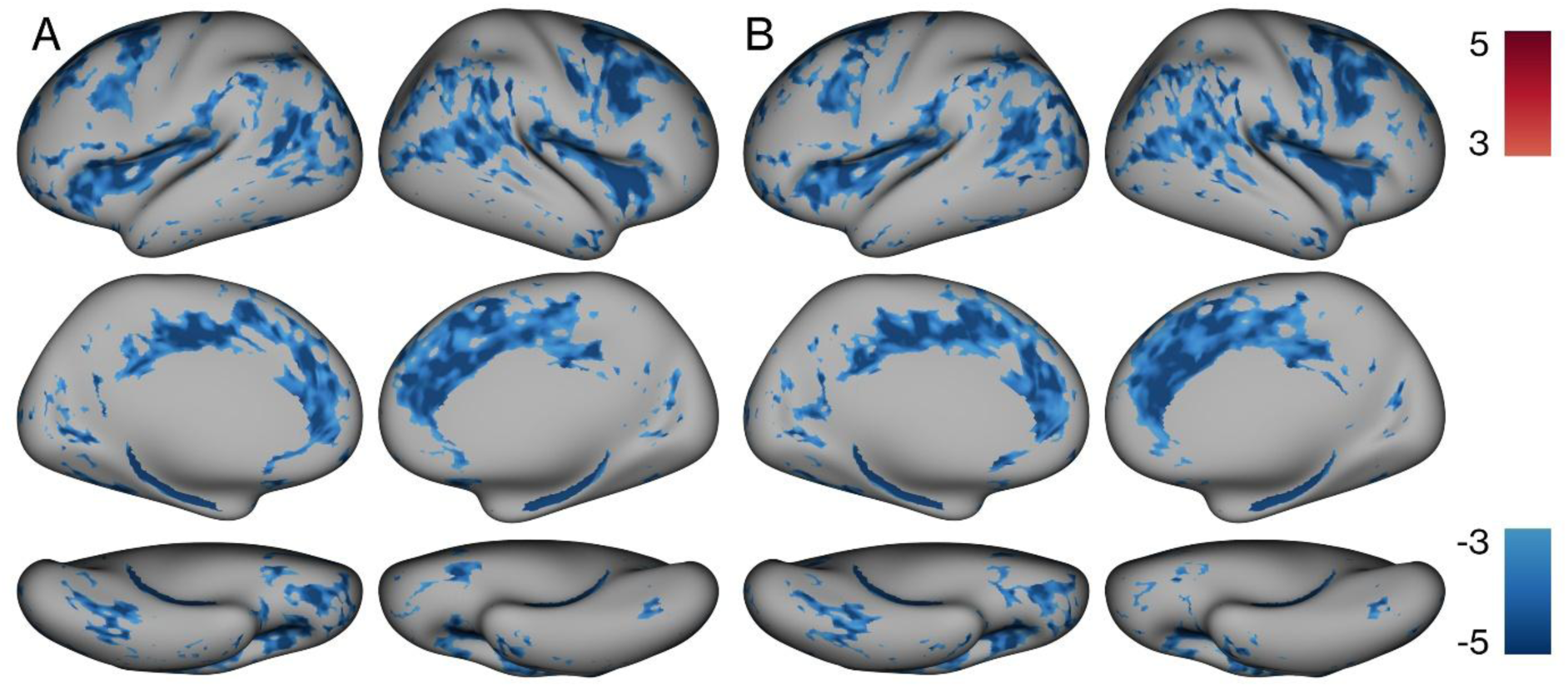
Thresholded statistical maps corresponding to vertex-wise interaction differences. Thresholded maps (*p* < 0.001) are shown for selected comparisons between pipelines, corresponding to the effect size maps presented in Figure 4. (A) LogHybrid vs FS6; (B) LogStrip vs FS6. The spatial distribution of significant differences is consistent with the observed effect size patterns, particularly in orbitofrontal regions. No spatially extended significant clusters were observed for the direct comparison between LogHybrid and LogStrip at this threshold. Thresholded maps (*p* < 0.001; |signed −log10(*p*)| ≥ 3) are shown. The color scale is fixed to ±5, and negative values indicate lower interaction values relative to the reference pipeline.

## Notes

### Competing Interest Statement

The authors have declared no competing interest.

